# The N764K and N856K mutations in SARS-CoV-2 Omicron BA.1 S protein generate potential cleavage sites for SKI-1/S1P protease

**DOI:** 10.1101/2022.01.21.477298

**Authors:** Halim Maaroufi

**Affiliations:** Institut de biologie intégrative et des systèmes (IBIS). Université Laval. Quebec, Qc G1V 0A6, Canada

**Keywords:** Spike protein, Omicron BA.1 variant, N764K/N856K, SKI-1/S1P (MBTPS1) protease

## Abstract

Spike (S) protein is a key protein in coronaviruses life cycle. SARS-CoV-2 Omicron BA.1 variant of concern (VoC) presents an exceptionally high number of 30 substitutions, 6 deletions and 3 insertions in the S protein. Recent works revealed major changes in the SARS-CoV-2 Omicron biological properties compared to earlier variants of concern (VoCs). Here, these major changes could be explained, at least in part, by the mutations N764K and/or N856K in S2 subunit. These mutations were not previously detected in other VoCs. N764K and N856K generate two potential cleavage sites for SKI-1/S1P serine protease, known to cleave viral envelope glycoproteins. The new sites where SKI-1/S1P could cleave S protein might impede the exposition of the internal fusion peptide for membrane fusion and syncytia formation. Based on the human protein atlas, SKI-1/S1P protease is not found in lung tissues (alveolar cells type I/II and endothelial cells), but present in bronchus and nasopharynx. This may explain why Omicron has change of tissue tropism. Viruses have evolved to use several host proteases for cleavage/activation of envelope glycoproteins. Mutations that allow viruses to change of protease may have a strong impact in host range, cell and tissue tropism, and pathogenesis.

## Introduction

The SARS-CoV-2 Omicron (B.1.1.529.1, BA.1) variant of concern (VoC) was first reported to WHO (Wealth Human organization) from South Africa on November 24^th^ of 2021. This VoC was from a specimen collected on November 9^th^ of 2021 in South Africa *(1, 2)*. Preliminary evidence suggests that Omicron variant increases the risk of reinfection and transmission compared to other variants, escapes from vaccines and is spreading rapidly across the world *(1)*. Indeed, Omicron has already been discovered in different countries around the world, and the number of nations reporting this variant continues to rise *(3)*. Omicron variant has an exceptionally high number of 30 substitutions, 6 deletions and 3 insertions in the S protein compared to the original Wuhan virus. Half of theses mutations are located in the receptor binding domain-RBD (amino acids 319-541), mutations N764K, D796Y, N856K, Q954H, N969K, L981F in S2 subunit were not previously detected in other VoCs, and mutations 69– 70del, T95I, G142D/143–145del, K417N, T478K, N501Y, N655Y, N679K, and P681H overlap with those in the alpha, beta, gamma, or delta variants *(4)*.

The SARS-CoV-2 S protein is a homotrimeric and multidomain. It is a key piece in coronaviruses cell life cycle: host range recognition (receptor-recognition), cell and tissue tropism, and pathogenesis *(5)*. Three categories of proteases cleave/activate S protein: type I transmembrane proprotein convertase (furin), cell surface type II transmembrane serine protease 2 (TMPRSS2, trypsin-like protease), and lysosomal cysteine proteases (cathepsin B/L). S protein is cleaved by Furin at residue 685 (^680^SPRRAR↓SV^687^) and generates S1 and S2 subunits. This cleavage at S1/S2 is important for efficient viral entry into cell. S1 subunit (residue 1–685) is divided into two domains, an N-terminal domain (NTD) and a C-terminal receptor-binding domain (RBD) that can function as viral receptors-binding *(6)*. S2 subunit (residue 686-1273) is multidomain and can be cleaved at residue 815 by TMPRSS2 producing S2’ (residue 816-1273), and thus allowing viral and host membranes fusion *(7)*.

Furin and SKI-1/S1P (subtilisin-kexin isozyme 1/site 1 protease) serine proteases are proprotein convertases related to subtilisin/Kexin (PCSKs) *(8)*. Furin (PCSK3) is important for efficient SARS-CoV-2 entry into cell. It cleaves/activates S protein at residue 685 and generates S1 and S2 subunits. Furin and SKI-1/S1P are part of subtilases that constitute the S8 family in clan SB of serine proteases (http://merops.sanger.ac.uk). Their active site is formed with a typical Asp, His and Ser catalytic triad and the oxyanion hole Asn. Furin (basic aa-specific) is able to cleave substrates at the [RK]-x-[RK]-↓-x consensus motif (x is any amino acid and ↓ is the cleavage site). Whereas, SKI-1/S1P (non-basic aa-specific) cleaves precursor proteins at the [RK]-x-[AILMFV]-[LTKF]-↓-x) consensus motif. Interestingly, amino acids around the cleavage site are important to determine the SKI-1/S1P activity in a specific subcellular compartment *(9)*. The membrane-bound SKI-1/S1P and Furin are the major ubiquitously expressed PCSKs in human tissues, and are used by viruses to produce mature virions activated for infection *(10, 11)*.

Here, the comparative analysis of Wuhan and Omicron S proteins revealed that the mutations N764K and N856K, not previously detected in other VoCs, generate new potential cleavage sites for SKI-1/S1P serine protease, known to cleave viral envelope glycoproteins. These potential cleavage sites for SKI-1/S1P serine protease could explain, at least in part, major biological changes in Omicron variant compared to Wuhan-Hu-1 and earlier VoCs.

## Results and discussion

Recently, Peacock and colleagues and Willett and coworkers demonstrated, *in vitro* and *ex vivo*, major changes in the SARS-CoV-2 Omicron biological properties *(12, 13)*. Indeed, compared to earlier SARS-CoV-2 VoCs, Omicron uses efficiently the endocytic pathway in a TMPRSS2-independent fashion, S protein has a decreased ability to induce cell syncytia formation, change in cell tropism and an expansion of ACE2 host range.

S protein is a key protein in coronaviruses life cycle. Amino acid mutation(s) of S protein at cleavage site(s) or mutation(s) that generate new cleavage site(s) may necessitate new proteases. Therefore, viruses have evolved to use several host proteases for cleavage/activation of envelope glycoproteins. For example, the mutation of the RRLA↓ motif in surface glycoprotein precursor (GP-C) to RRRR↓ does allow/constraint arenavirus lymphocytic choriomeningitis virus (LCMV) to use Furin instead of SKI-1/S1P protease *(14)*. Change of proteases that process viral envelope glycoproteins may have a strong impact in host range, cell and tissue tropism, and pathogenesis *(15, 16)*.

### The mutations N764K and N856K generate potential cleavage sites for SKI-1/S1P protease

Major changes in biological properties of SARS-CoV-2 Omicron BA.1 variant compared to Wuhan-Hu-1 and earlier VoCs could be explained, at least in part, by the mutations N764K and/or N856K in S protein. These mutations were not previously detected in other VoCs. N764K and N856K generate new potential cleavage sites ^764^KRAL^767^ ↓TG and ^856^KGLT^859^↓VL (Fig. 1 and Fig. 2A) for SKI-1/S1P serine protease (UniProt ID: Q14703), known to cleave viral envelope glycoproteins *(17, 18)*. Interestingly, one part of ^764^KRAL^767^↓TG motif is corresponding to a Furin (-like) cleavage site ^764^**KR**↓A**L**^767^. The latter is similar to ^814^**KR**↓S**F**^817^ that is cleaved by the single Arg-specific enzyme TMPRSS2 *(19)*.

**Fig. 1.**
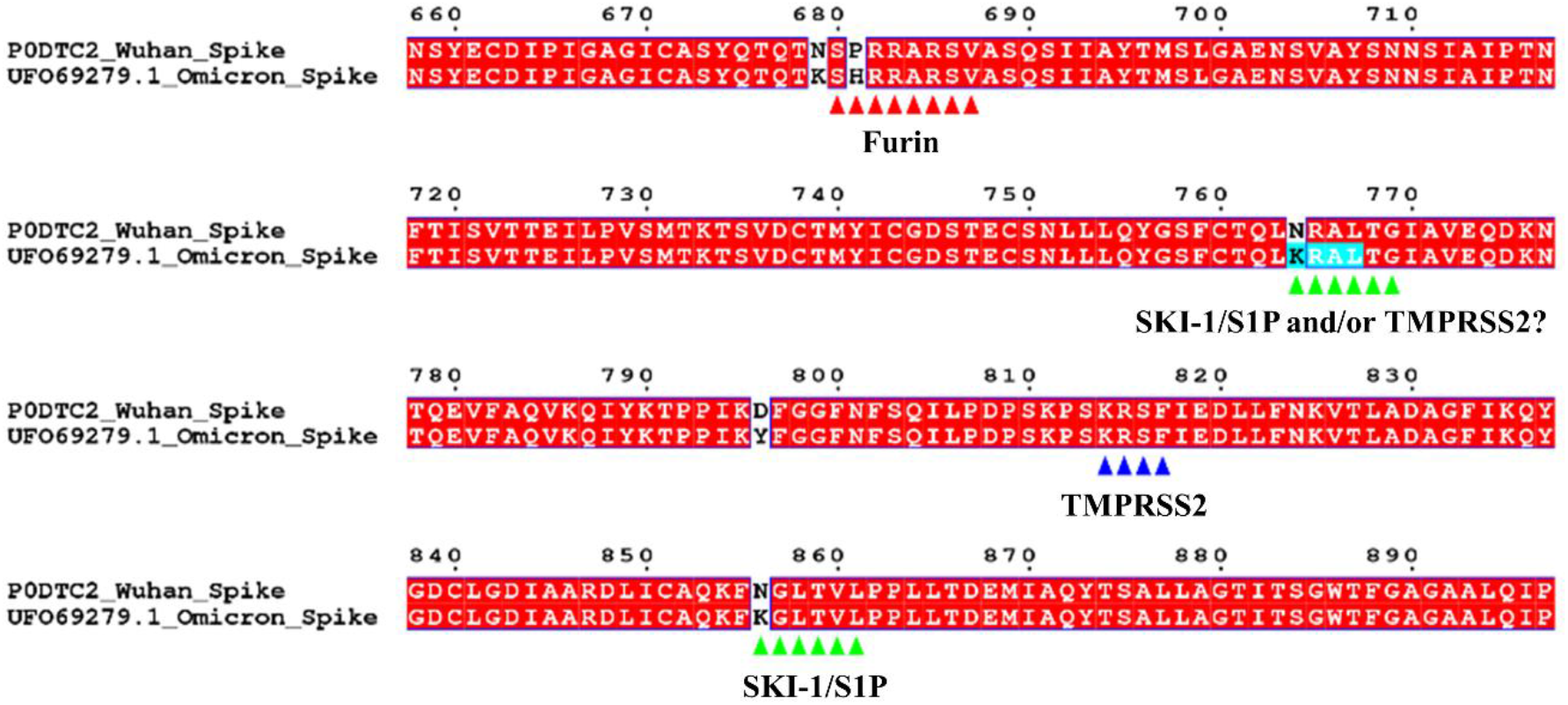
Cleavage sites on SARS-CoV-2 S protein. Wuhan S protein cleavage sites of Furin (S1/S2) (^680^SPRRAR↓SV^687^, red arrows head) and TMPRSS2 (S2’) (^814^KR↓SF^817^, blue arrows head). Potential Omicron BA.1 variant S protein cleavage sites of SKI-1/S1P protease (^764^KRAL^767^ ↓TG and ^856^KGLT^859^↓VL, green arrows head) and potential TMPRSS2 (^764^KR↓AL^767^, highlighted in cyan). The figure was prepared with ESPript (http://espript.ibcp.fr).

**Fig. 2.**
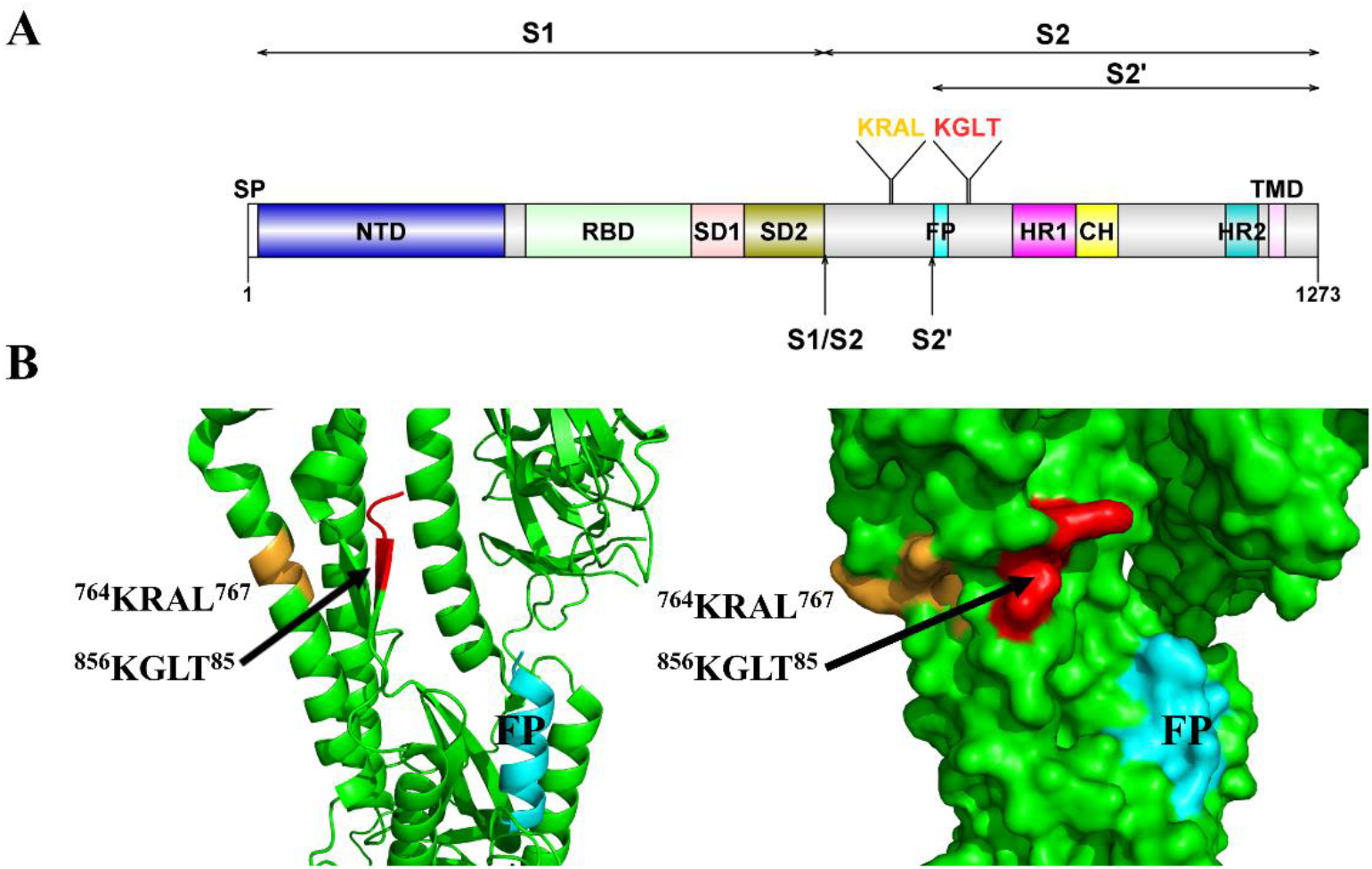
The potential cleavage sites for SKI-1/S1P protease in the SARS-CoV-2 Omicron BA.1 S protein. **(A)** Diagram representation of S protein (S1 subunit (residue 1–685), S2 subunit (residue 686-1273), S2’ (residue 816-1273)) colored by domain. Signal peptide (SP), N-terminal domain (NTD), receptor-binding domain (RBD), subdomains 1 and 2 (SD1 and SD2), S1/S2 Furin cleavage site, S2’ TMPRSS2 cleavage, fusion peptide (FP), heptad repeat 1 and 2 (HR1 and HR2), central helix (CH), transmembrane domain (TMD), and the localization of potential cleavage sites ^764^KRAL^767^ and ^856^KGLT^859^ For SKI-1/S1P protease. **(B)** Structure of Omicron S protein monomer (PDB ID: 7T9J_C) in cartoon (left) and surface (right), with potential cleavage sites^764^KRAL^767^ (in orange) and ^856^KGLT^859^ (in red) of PCSK_SKI-1 protease, and fusion peptide (FP, in cyan). Of note, the region between amino acids 828 and 852, close to ^856^KGLT^859^, is not resolved in all structures of SARS-CoV-2 S proteins. The figure 1A and 1B were prepared with DOG (http://dog.biocuckoo.org/down.php) and PyMol (www.pymol.org), respectively.

^764^KRAL^767^ ↓TG and ^856^KGLT^859^↓VL are solvent exposed what is necessary for efficient proteolysis (Fig. 2B, right). In addition, ^856^KGLT^859^↓VL is found near a region, between amino acids 828 and 852 (Fig. 2B, left), not resolved in all structures of SARS-CoV-2 S proteins, and hence considered intrinsically disordered (https://disprot.org/DP02772). It has been reported that cleavage sites in Zika virus tend to occur in or at least in close proximity to the regions of substrates with increased flexibility and/or intrinsic disorder *(20)*. This suggests that ^856^KGLT^859^↓VL site is more likely to be cleaved by SKI-1/S1P.

### SKI-1/S1P is a key player in the cell

SKI-1/S1P secretory serine protease, also named membrane-bound transcription factor site-1 protease (MBTPS1) cleaves precursor proteins at ([RK]-x-[AILMFV]-[LTKF]-↓-x) site (x is any amino acid and ↓ is the cleavage site). Interestingly, amino acids around the cleavage site are important to determine the SKI-1/S1P activity in a specific subcellular compartment *(9)*. SKI-1/S1P and Furin are the major ubiquitously expressed PCSKs. They are used by several viruses to activate their proteins for infection *(10, 11)*. SKI-1/S1P localizes in endoplasmic reticulum membrane, Golgi apparatus membrane, and may sort to other organelles, including lysosomal and/or endosomal compartments *(10)*. It is supposed to be fully active in the cis/medial Golgi. In cell, SKI-1/S1P is responsible of the activation of several transcription factors. Indeed, SKI-1/S1P plays a major role in cholesterol metabolism because it cleaves/activates sterol regulatory element binding proteins (SREBP-1 and SREBP-2) that modulates the expression of genes implicated in cholesterol synthesis *(21)*. It has been observed that the hyperinflammation in COVID-19 patients is induced by the alteration of cholesterol and lipid synthesis *(22)*. In addition, Lee and collaborators reported that COVID-19 patients’ blood contains an SREBP-2 C-terminal fragment (indicator for disease severity), likely due to an upregulation of the SKI-1/S1P by SARS-CoV-2 Wuhan-Hu-1*(23)*. It is tempting to speculate that Omicron by hijacking SKI-1/S1P for cleaving S protein might not over activate SREBP-2. This may explain, at least in part, why Omicron variant causes mild disease. Another important role is played by SKI-1/S1P protease in the generation of antibody secreting cells and reprogramming for secretory activity. Indeed, the inhibition of SKI-1/S1P by PF-429242 (a reversible, competitive aminopyrrolidineamide inhibitor) induces a pronounced reduction in the ability of plasmablasts CD38^low^ populations to secret IgG compared to IgM plasmablasts *(24)*. All these facts let suppose that SARS-CoV-2 by manipulating host SKI-1/S1P protease, a key player in cell, disrupt cell homeostasis, and hence induces COVID-19.

### Potential role of SKI-1/S1P in change of Omicron biological properties

In Omicron variant, the impairment of cell surface entry and syncytia formation may be due to the introduction of ^764^KRAL^767^ ↓TG and ^856^KGLT^859^↓VL SKI-1/S1P cleavage sites that may produce at least an S2’’ subunit (residue 768-1273) and/or S2’’’ (residue 860-1273) without exposition of the internal fusion peptide for membrane fusion (Fig. 1 and Fig. 2A). Interestingly, based on the human protein atlas (https://www.proteinatlas.org/ENSG00000140943-MBTPS1/tissue), SKI-1/S1P protein (gene name: MBTPS1) is found in macrophages of lung but not in lung tissues (alveolar cells type I/II and endothelial cells), whereas, it is present in bronchus (respiratory epithelial cells) and nasopharynx (respiratory epithelial cells). This could explain why the replication of Omicron is reduced in lower respiratory tissues, but possibly increased in the upper respiratory tract *(12, 13, 25)*. These findings are in accord with works that reported that modulation of cleavage site in S protein appears to be correlated with changes in cell and tissue tropism and pathogenicity *(26, 27)*.

### SKI-1/S1P contains, in addition to catalytic domain, two other domains

An experimental structure of SKI-1/S1P is not yet available. Thanks to AlphaFold 2 (AF2) to have predicted 3D model of SKI-1/S1P (https://alphafold.ebi.ac.uk/entry/Q14703) with very high confidence for molecular docking. AF2 model revealed that SKI-1/S1P is formed by three domains: a catalytic domain (CD) with a typical Asp218, His249 and Ser414 catalytic triad and the oxyanion hole Asn338, P domain (PD) that apparently stabilizes the catalytic pocket, and an all beta fold domain (Ig-likeD) (Fig. 3A). Electrostatic potential surface representation of CD revealed four subsites, S1/S2 in active site and S3/S4 distal from the catalytic triad (Fig. 3C). Structural similarity search (Fig. 1S) showed that PD is similar to intraflagellar transport (IFT) 52N (IFT52N) domain (PDB ID: 5FMR_B). IFT52N is a subunit of IFT complex involved in the assembly and maintenance of eukaryotic cilia *(28)*. The epithelium in airways takes an important role in the defense against pathogens *(29)*. Human coronaviruses were shown to target ciliated cells *(30, 31)*, revealing their role in coronavirus pathogenesis. Whereas, all beta fold domain presents structural similarity with immunoglobulin-like (Ig-like) domain (PDB ID: 2E6J_A) (Fig. S1). Ig-like domains are involved in protein-protein interactions, often with other Ig-like domains by their beta-sheets *(32)*. Ig-like domain is present in hydrocephalus-inducing protein homolog (hydin) that is required for ciliary motility (UniProt ID: Q4G0P3).

**Fig. 3.**
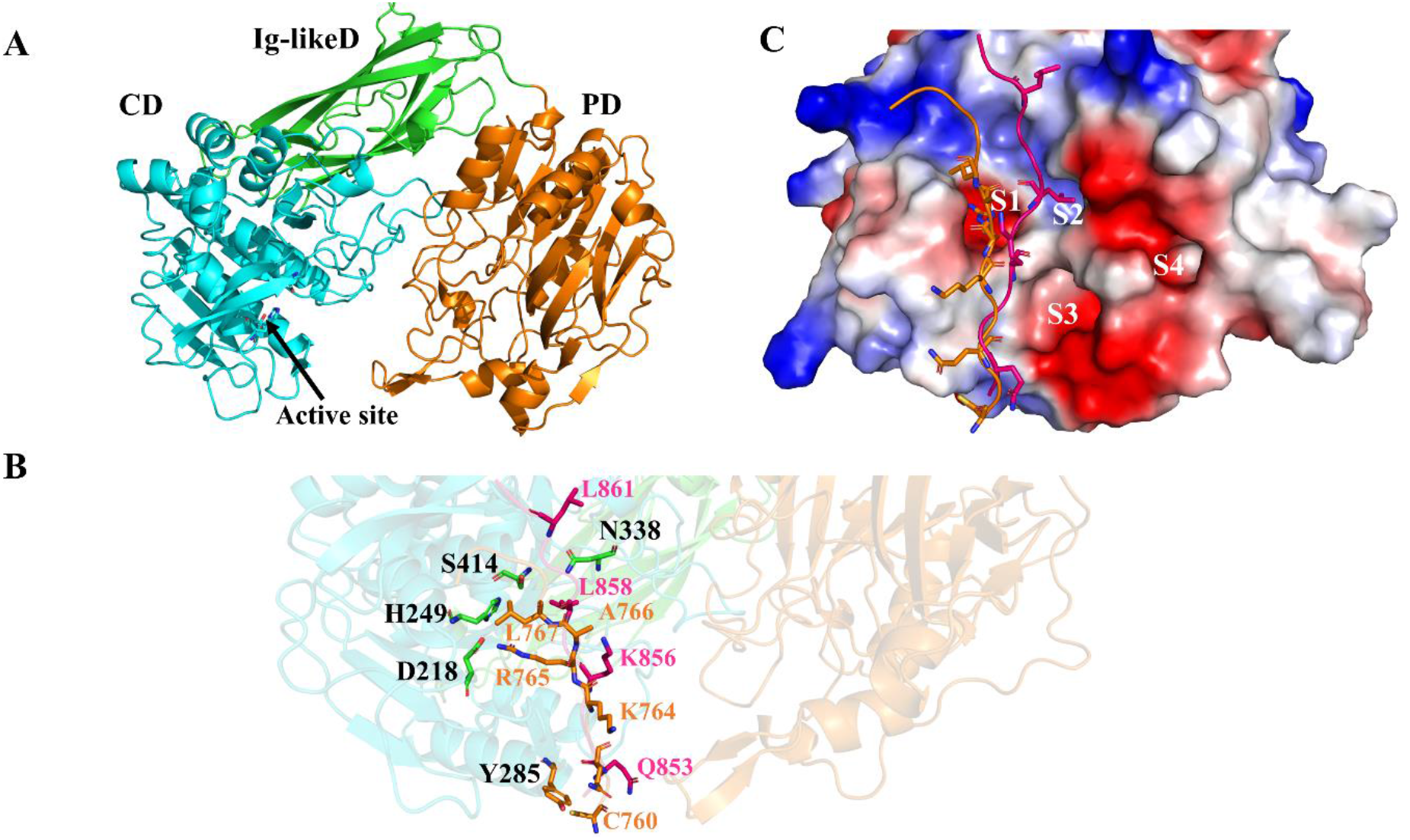
The structure of SKI-1/S1P predicted by AF2 with docked Omicron S peptides. **(A)** SKI-1/S1P structure with three domains catalytic domain (CD), P domain (PD) and immunoglobulin-like domain (Ig-likeD). The active site in CD (arrow). **(B)** Peptides containing SKI-1/S1P cleavage sites, CTQL^764^**KRAL**^767^TGIA (in orange) and AQKF^856^**KGLT**^859^VLPP (in pink), by blind docking are localized in catalytic domain. Active site is formed by Asp218, His249 and Ser414 catalytic triad and the oxyanion hole Asn338. Amino acid Tyr285 may play a role in the molecular recognition of the peptide. For bonds between SKI-1/S1P and peptides, see Tables S1-2. **(C)** Electrostatic potential surface representation of CD with docked peptides. Subsites formed on the face of the CD, S1/S2 in active site and S3/S4 distal from the catalytic triad.

### Molecular docking of peptides from S protein into SKI-1/S1P

Blind docking of Omicron S protein peptides ^760^CTQL**<u>K</u>**RA**<u>L</u>**TGIA^771^ (peptide764) and ^852^AQKF**<u>K</u>**GL**<u>T</u>**VLPP^863^ (peptide856) into SKI-1/S1P (Fig. 3B) showed that peptide764 establishes more hydrogen bonds with SKI-1/S1P than peptide856 (Table S1 and S2). In addition, peptide764 forms salt bridges between Asp218 and Arg765. Predicted ΔG (binding free energy) of SKI-1/S1P-peptide764 complex is −58.98 kcal mol^−1^ in which Arg765, Leu763 and Leu767 participate with −10.43, −6.32 and −5.05 kcal mol^−1^, respectively (Table S3). Whereas, predicted ΔG of SKI-1/S1P-peptide856 is −30.00 kcal mol^−1^ in which Phe855, Val560 and Leu858 participate with −6.45,−5.56 and −5.26 kcal mol^−1^, respectively (Table S4). These molecular docking results indicate that SKI-1/S1P probably has more affinity for the peptide764, suggesting that SKI-1/S1P may cleave Omicron S protein better in the site 764 than 856. In addition, during the writing of this manuscript two sub-lineages BA.2 and BA.3 of Omicron have been discovered. Interestingly, BA.2 and BA.3 do not have the mutation N856K, thus no potential cleavage site ^856^KGLT^859^↓VL for SKI-1/S1P serine protease. The first medical observations about BA.2 lineage suggest there is no great difference in disease severity compared to BA.1. These observations and molecular docking results suggest that SKI-1/S1P probably cleave ^764^KRAL^767^ ↓TG site in BA.1, BA.2 and BA.3 lineages.

Finally, should an experimental assessment of these findings show that SKI-1/S1P protease cleaves Omicron S protein as in other enveloped RNA viruses? Thus, SKI-1/S1P will be a relevant target to impede Omicron infection. In fact, the inhibition of SKI-1/S1P has already been tested in coronaviruses. Plegge and colleagues reported that SKI-1/S1P inhibitor PF-429242 inhibited MERS-CoV and SARS-CoV S protein-driven entry efficiently by blocking processing of the S proteins *(33, 34)*. Furthermore, PF-429242 showed strong dose-dependent reduction of SARS-CoV-2 (USA/WA-1/2020 strain) replication with cytotoxicity only at high concentration *(35)*.

## Material and methods

### Sequence analysis

To see the impact of the mutations in Omicron S protein on short linear motifs (SLiMs), SARS-CoV-2 S protein sequences of Wuhan (UniProt ID: P0DTC2) and Omicron variant (GenBank ID: UFO69279.1) from Belgium was scanned with the eukaryotic linear motif (ELM) resource (http://elm.eu.org/).

### 3D modeling and molecular docking

The 3D model of human SKI-1/S1P predicted by AlphaFold 2 (AF2) was downloaded from https://alphafold.ebi.ac.uk/entry/Q14703 *(36)*. It was used in molecular dockings. The region between amino acids 828 and 856, close to ^856^KGLT^859^, is not resolved in the structure of Omicron S protein (PDB ID: 7T9J_C). Therefore, to have the structure of ^852^AQKF**<u>K</u>**GL**<u>T</u>**VLPP^863^peptide, the 3D model of Omicron S was predicted by AF2 software download at home *(37)*.

Before starting blind peptide-protein docking, the validation of the accuracy of HPEPDOCK docking software was conducted by docking the subunits of crystal structure of the complex tankyrase-2 RD-human peptide SH3BP2 (PDB ID: 3TWR) *(38)*. Thus, in docking protocol, the coordinates of each separated molecules were used as ligand (PDB ID: 3TWR _G) and receptor (PDB ID: 3TWR _C) for SH3BP2 peptide (LPHLQ**RSPPDG**QSFRS) and tankyrase-2, respectively. Indeed, HPEPDOCK was able to produce a similar docking pose for each control protein with respect to its biological conformation in the co-crystallised protein-peptide complex.

The potential interactions of SARS-CoV-2 Omicron S protein with the human SKI-1/S1P protease were tested by molecular docking. Indeed, the 3D coordinates of ^760^CTQL**<u>K</u>**RA**<u>L</u>**TGIA^771^ (peptide764) and ^852^AQKF**<u>K</u>**GL**<u>T</u>**VLPP^863^ (peptide856) peptides were extracted from Omicron S protein structure (PDB ID: 7T9J_C) and 3D model of Omicron S predicted by AF2, respectively. These peptides were docked into the 3D model of human SKI-1/S1P using HPEPDOCK software *(38)*.

The predicted binding free energy of protein-peptide complex was determined by MM/GBSA. The MM/GBSA is calculated based on the ff02 force field, the implicit solvent model and the GB^OBC1^ model (interior dielectric constant = 1). The whole system was minimized for 5000 steps with a cutoff distance of 12 Å for van der Waals interactions (2000 cycles of steepest descent and 3000 cycles of conjugate gradient minimizations) *(39)*.

Structural similarity of the predicted SKI-1/S1P structure was performed with PDBeFold (https://www.ebi.ac.uk/msd-srv/ssm/ssmstart.html) and http://ekhidna2.biocenter.helsinki.fi/dali/ against PDB database and human AlphaFold database (https://alphafold.ebi.ac.uk/). The electrostatic potential surfaces of 3D models and images were generated with PyMOL software (http://pymol.org/).

## Acknowledgments

I would like to thank the IBIS bioinformatics group for their help.

## Supplementary material

**Table S1.** Predicted bonds between SKI-1/S1P and peptide764 (^760^CTQLKRALTGIA^771^)

| Hydrogen bonds |  |  |  |
| --- | --- | --- | --- |
| ## | SKI-1/S1P | Dist. [Å] | Peptide764 |
| 1 | THR 286[ N ] | 3.62 | THR 761[ O ] |
| 2 | THR 286[ OG1] | 2.62 | THR 761[ OG1] |
| 3 | SER 284[ N ] | 3.08 | LEU 763[ O ] |
| 4 | ARG 408[ NH1] | 2.73 | ILE 770[ O ] |
| 5 | TYR 285[ OH ] | 3.59 | GLN 762[ NE2] |
| 6 | SER 284[ O ] | 3.01 | LEU 763[ N ] |
| 7 | GLN 282[ O ] | 3.29 | ARG 765[ N ] |
| 8 | ASP 218*[ OD2] | 3.24 | ARG 765[ NH1] |
| 9 | ASP 218*[ OD1] | 2.66 | ARG 765[ NH2] |
| 10 | THR 219[ OG1] | 3.87 | ARG 765[ NH2] |
| 11 | GLN 282[ OE1] | 3.07 | LEU 767[ N ] |
| Salt bridges |  |  |  |
| 1 | ASP 218*[ OD1] | 3.49 | ARG 765[ NH1] |
| 2 | ASP 218*[ OD2] | 3.24 | ARG 765[ NH1] |
| 3 | ASP 218*[ OD1] | 2.66 | ARG 765 NH2] |
| 4 | ASP 218*[ OD2] | 3.71 | ARG 765[ NH2] |

**N.B.** \* Asp218, His249 and Ser414 catalytic triad

**Table S2.** Predicted bonds between SKI-1/S1P and peptide856 (^852^AQKFKGLTVLPP^863^)

| Hydrogen bonds |  |  |  |
| --- | --- | --- | --- |
| ## | SKI-1/S1P | Dist. [Å] | Peptide856 |
| 1 | THR 286[ N ] | 3.87 | GLN 853[ O ] |
| 2 | SER 411[ OG ] | 3.75 | PRO 862[ O ] |
| 3 | SER 287[ OG ] | 3.81 | ALA 852[ N ] |
| 4 | SER 284[ O ] | 3.55 | PHE 855[ N ] |
| 5 | SER 411[ O ] | 2.66 | LEU 861[ N ] |

**Table S3.** Predicted binding Free Energy (ΔG) of residue from peptide764 (^760^CTQL**KRAL**TGIA^771^) and twelve high ΔG residues of SKI-1/S1P.

| Residue (peptide764) | $\Delta G$ (kcal mol<sup>-1</sup>) | Residue (SKI-1/S1P) | $\Delta G$ (kcal mol<sup>-1</sup>) |
| --- | --- | --- | --- |
| ARG-765 | -10.43 | ASP-218* | -6.29 |
| LEU-763 | -6.32 | TYR-285 | -4.89 |
| LEU-767 | -5.05 | VAL-283 | -3.43 |
| ILE-760 | -3.73 | SER-284 | -3.39 |
| GLN-762 | -2.54 | PHE-278 | -3.26 |
| LYS-764 | -1.42 | GLN-282 | -3.15 |
| THR-761 | -0.91 | HIS-249* | -1.65 |
| ALA766 | -0.46 | LEU-410 | -1.63 |
| GLY-769 | 0.31 | ILE-308 | -1.5 |
| ALA771 | 0.55 | ARG-408 | -1.38 |
| THR-768 | 0.88 | THR-286 | -1.26 |
| CYS-760 | 1.15 | LEU-247 | -1.09 |

**N.B.** $\Delta G$ of SKI-1/S1P-peptide764 complex is -58.98 kcal mol<sup>-1</sup>
\* Asp218, His249 and Ser414 catalytic triad

**Table S4.**
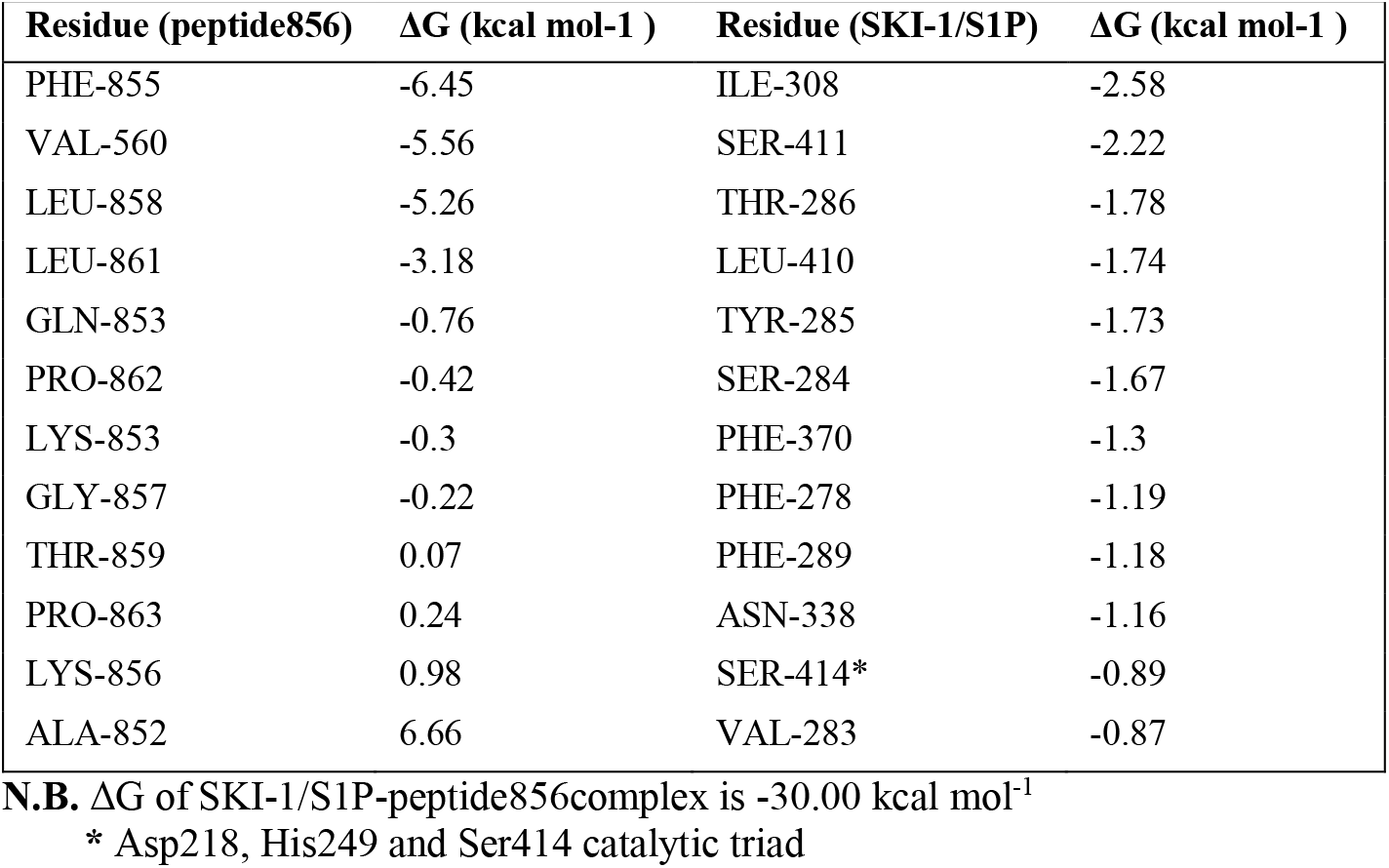
Predicted binding Free Energy (ΔG) of residue from peptide856 (^852^AQKF**KGLT**VLPP^863^) and twelve high ΔG residues of SKI-1/S1P.

| Residue (peptide856) | $\Delta G$ (kcal mol<sup>-1</sup>) | Residue (SKI-1/S1P) | $\Delta G$ (kcal mol<sup>-1</sup>) |
| --- | --- | --- | --- |
| PHE-855 | -6.45 | ILE-308 | -2.58 |
| VAL-560 | -5.56 | SER-411 | -2.22 |
| LEU-858 | -5.26 | THR-286 | -1.78 |
| LEU-861 | -3.18 | LEU-410 | -1.74 |
| GLN-853 | -0.76 | TYR-285 | -1.73 |
| PRO-862 | -0.42 | SER-284 | -1.67 |
| LYS-853 | -0.3 | PHE-370 | -1.3 |
| GLY-857 | -0.22 | PHE-278 | -1.19 |
| THR-859 | 0.07 | PHE-289 | -1.18 |
| PRO-863 | 0.24 | ASN-338 | -1.16 |
| LYS-856 | 0.98 | SER-414* | -0.89 |
| ALA-852 | 6.66 | VAL-283 | -0.87 |

**N.B.** $\Delta G$ of SKI-1/S1P-peptide856complex is -30.00 kcal mol<sup>-1</sup>
\* Asp218, His249 and Ser414 catalytic triad

**Fig. S1.**
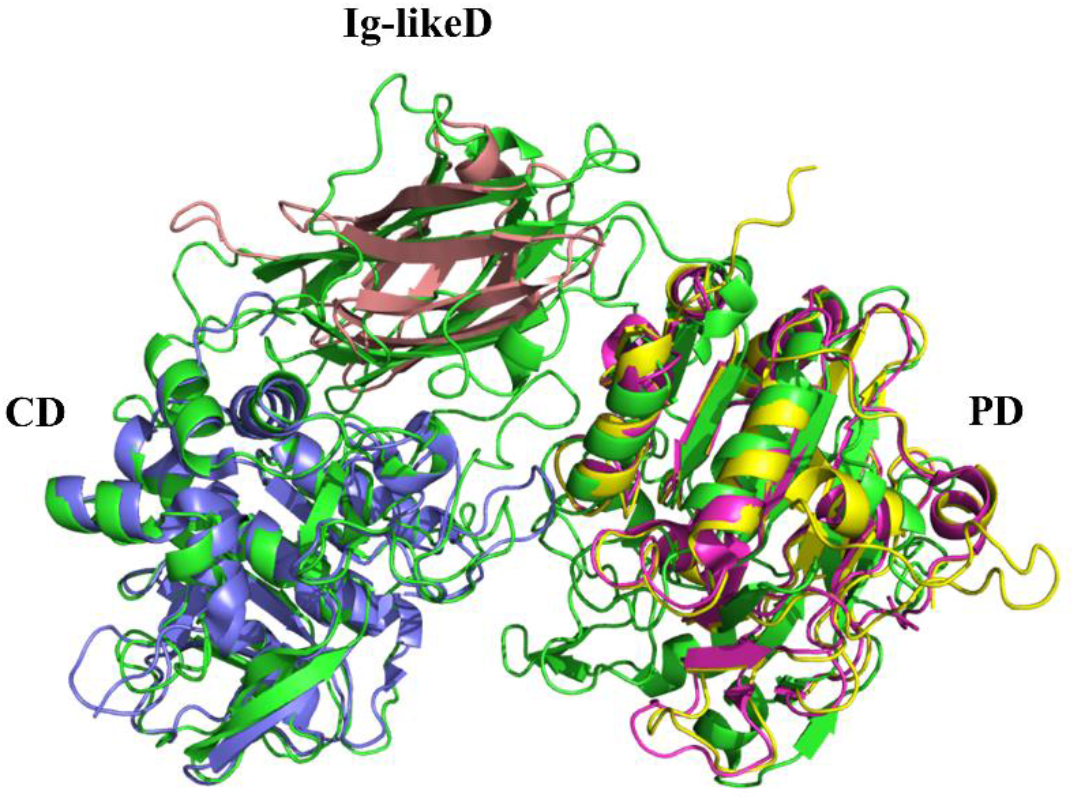
Superposition of domains similar to those of SKI-1/S1P. Catalytic domain (CD) of SKI-1/S1P (in green) superposes with Subtilisin BPN’ of *Bacillus amyloliquefaciens* (in blue, PDB ID: 1LW6_A). All beta domain (Ig-likeD in green) superposes with immunoglobulin-like domain of C-terminal PapD-like domain from human hydin protein (in salmon, PDB ID: 2E6J_A). The P domain (PD in green) superposes with *Chlamydomonas reinhardtii* intraflagellar transport 52N (IFT52N) domain (in pink, PDB ID: 5FMR_B) and human IFT52 (in yellow, UniProt ID: Q9Y366; AF2 3D model).

## References

1. Classification of Omicrom (B.1.1.529): SARS-CoV-2 Variant of Concern. https://www.who.int/news/item/26-11-2021-classification-of-omicron-(b.1.1.529)-sars-cov-2-variant-of-concern

2. K. Kupferschmidt, Startling new variant raises urgent questions. Science 374, 1178–80 (2021).

3. European Centre for Disease Prevention and Control. Implications of the emergence and spread of the SARSCoV-2 B.1.1.529 variant of concern (Omicron), for the EU/EEA. 26 November 2021. ECDC: Stockholm; 2021.

4. GISAID. Tracking of variants. 2021. https://www.gisaid.org/hcov19-variants/ (accessed Nov 30, 2021)

5. F. Li, Receptor recognition mechanisms of coronaviruses: a decade of structural studies. J Virol. 89, 1954–1964 (2015).

6. F. Li, Evidence for a common evolutionary origin of coronavirus spike protein receptor-binding subunits. J Virol. 86, 2856–2858 (2012).

7. E. Paszti-Gere, R. F. Barna, C. Kovago, I. Szauder, G. Ujhelyi, C. Jakab, N. Meggyesházi, A. Szekacs, Changes in the distribution of type II transmembrane serine protease, TMPRSS2 and in paracellular permeability in IPEC-J2 cells exposed to oxidative stress. Inflammation. 38, 775–83 (2015).

8. N. G. Seidah, A. Prat, The biology and therapeutic targeting of the proprotein convertases. Nat Rev Drug Discov. 11, 367–83 (2012).

9. D. J. Burri, G. Pasqual, C. Rochat, N. G. Seidah, A. Pasquato, S. Kunz, Molecular characterization of the processing of arenavirus envelope glycoprotein precursors by subtilisin kexin isozyme-1/site-1 protease. J Virol. 86, 4935–46 (2012).

10. N. G. Seidah, S. J. Mowla, J. Hamelin, A. M. Mamarbachi, S. Benjannet, B. B. Touré, A. Basak, J. S. Munzer, J. Marcinkiewicz, M. Zhong, J. C. Barale, C. Lazure, R. A. Murphy, M. Chrétien, M. Marcinkiewicz, Mammalian subtilisin/kexin isozyme SKI-1: A widely expressed proprotein convertase with a unique cleavage specificity and cellular localization. Proc Natl Acad Sci U S A. 96, 1321–6 (1999).

11. W. J. Van de Ven, J. W. Creemers, A. J. Roebroek, Furin: the prototype mammalian subtilisin-like proprotein-processing enzyme. Endoproteolytic cleavage at paired basic residues of proproteins of the eukaryotic secretory pathway. Enzyme. 45, 257–70 (1991).

12. T. P. Peacock et al., The SARS-CoV-2 variant, Omicron, shows rapid replication in human primary nasal epithelial cultures and efficiently uses the endosomal route of entry. bioRxiv, 2021.2012.2031.474653 (2022).

13. B. J. Willett et al., The hyper-transmissible SARS-CoV-2 Omicron variant exhibits significant antigenic change, vaccine escape and a switch in cell entry mechanism. medRxiv, 2022.2001.2003.21268111 (2022).

14. J. M. Rojek, G. Pasqual, A. B. Sanchez, N. T. Nguyen, J. C. de la Torre, S. Kunz, Targeting the proteolytic processing of the viral glycoprotein precursor is a promising novel antiviral strategy against arenaviruses. J Virol. 84, 573–84 (2010).

15. H. D. Klenk, W. Garten, Host cell proteases controlling virus pathogenicity. Trends Microbiol. 2, 39–43 (1994).

16. Y. Nagai, Protease-dependent virus tropism and pathogenicity. Trends Microbiol. 1, 81–7 (1993).

17. O. Lenz, J. ter Meulen, H. D. Klenk, N. G. Seidah, W. Garten, The Lassa virus glycoprotein precursor GP-C is proteolytically processed by subtilase SKI-1/S1P. Proc Natl Acad Sci U S A. 98, 12701–5 (2001).

18. M. J. Vincent, A. J. Sanchez, B. R. Erickson, A. Basak, M. Chretien, N. G. Seidah, S. T. Nichol, Crimean-Congo hemorrhagic fever virus glycoprotein proteolytic processing by subtilase SKI-1. J Virol. 77, 8640–9 (2003).

19. B. F. Hörnich, A. K. Großkopf, S. Schlagowski, M. Tenbusch, H. Kleine-Weber, F. Neipel, C. Stahl-Hennig, A. S. Hahn, SARS-CoV-2 and SARS-CoV Spike-Mediated Cell-Cell Fusion Differ in Their Requirements for Receptor Expression and Proteolytic Activation. J Virol. 95(2021), doi:10.1128/jvi.00002-21.

20. R. Giri, D. Kumar, N. Sharma, V. N. Uversky, Intrinsically Disordered Side of the Zika Virus Proteome. Front Cell Infect Microbiol. 6, 144 (2016).

21. J. Sakai, R. B. Rawson, P. J. Espenshade, D. Cheng, A. C. Seegmiller, J. L. Goldstein, M. S. Brown, Molecular identification of the sterol-regulated luminal protease that cleaves SREBPs and controls lipid composition of animal cells. Mol Cell. 2, 505–14 (1998).

22. Y. Sohrabi, H. Reinecke, R. Godfrey, Altered Cholesterol and Lipid Synthesis Mediates Hyperinflammation in COVID-19. Trends Endocrinol Metab. 32, 132–134 (2021).

23. W. Lee, J. H. Ahn, H. H. Park, H. N. Kim, H. Kim, Y. Yoo, H. Shin, K. S. Hong, J. G. Jang, C. G. Park, E. Y. Choi, J. S. Bae, Y. K. Seo, COVID-19-activated SREBP2 disturbs cholesterol biosynthesis and leads to cytokine storm. Signal Transduct Target Ther. 5, 186 (2020).

24. M. Al-Maskari, M. A. Care, E. Robinson, M. Cocco, R. M. Tooze, G. M. Doody, Site-1 protease function is essential for the generation of antibody secreting cells and reprogramming for secretory activity. Sci Rep. 8, 14338 (2018).

25. R. Abdelnabi et al., The omicron (B.1.1.529) SARS-CoV-2 variant of concern does not readily infect Syrian hamsters. bioRxiv, 2021.2012.2024.474086 (2021).

26. J. K. Millet, G. R. Whittaker, Host cell proteases: Critical determinants of coronavirus tropism and pathogenesis. Virus Res. 202, 120–134 (2015).

27. F. P. Tay, M. Huang, L. Wang, Y. Yamada, D. X. Liu, Characterization of cellular furin content as a potential factor determining the susceptibility of cultured human and animal cells to coronavirus infectious bronchitis virus infection. Virology. 433, 421–30 (2012).

28. M. Taschner, K. Weber, A. Mourão, M. Vetter, M. Awasthi, M. Stiegler, S. Bhogaraju, E. Lorentzen, Intraflagellar transport proteins 172, 80, 57, 54, 38, and 20 form a stable tubulin-binding IFT-B2 complex. Embo j. 35, 773–90 (2016).

29. A. E. Tilley, M. S. Walters, R. Shaykhiev, R. G. Crystal, Cilia dysfunction in lung disease. Annu Rev Physiol. 77, 379–406 (2015).

30. N. Zhu, W. Wang, Z. Liu, C. Liang, F. Ye, B. Huang, L. Zhao, H. Wang, W. Zhou, Y. Deng, L. Mao, C. Su, G. Qiang, T. Jiang, J. Zhao, G. Wu, J. Song, W. Tan, Morphogenesis and cytopathic effect of SARS-CoV-2 infection in human airway epithelial cells. Nat Commun. 11, 3910 (2020).

31. R. Robinot, M. Hubert, G. D. de Melo, F. Lazarini, T. Bruel, N. Smith, S. Levallois, F. Larrous, J. Fernandes, S. Gellenoncourt, S. Rigaud, O. Gorgette, C. Thouvenot, C. Trébeau, A. Mallet, G. Duménil, S. Gobaa, R. Etournay, P. M. Lledo, M. Lecuit, H. Bourhy, D. Duffy, V. Michel, O. Schwartz, L. A. Chakrabarti, SARS-CoV-2 infection induces the dedifferentiation of multiciliated cells and impairs mucociliary clearance. Nat Commun. 12, 4354 (2021).

32. D. M. Halaby, A. Poupon, J. Mornon, The immunoglobulin fold family: sequence analysis and 3D structure comparisons. Protein Eng. 12, 563–71 (1999).

33. T. Plegge, M. Spiegel, N. Krüger, I. Nehlmeier, M. Winkler, M. González Hernández, S. Pöhlmann, Inhibitors of signal peptide peptidase and subtilisin/kexin-isozyme 1 inhibit Ebola virus glycoprotein-driven cell entry by interfering with activity and cellular localization of endosomal cathepsins. PLoS One. 14, e0214968 (2019).

34. J. L. Hawkins, M. D. Robbins, L. C. Warren, D. Xia, S. F. Petras, J. J. Valentine, A. H. Varghese, I. K. Wang, T. A. Subashi, L. D. Shelly, B. A. Hay, K. T. Landschulz, K. F. Geoghegan, H. J. Harwood, Pharmacologic inhibition of site 1 protease activity inhibits sterol regulatory element-binding protein processing and reduces lipogenic enzyme gene expression and lipid synthesis in cultured cells and experimental animals. J Pharmacol Exp Ther. 326, 801–8 (2008).

35. R. Wang, C. R. Simoneau, J. Kulsuptrakul, M. Bouhaddou, K. A. Travisano, J. M. Hayashi, J. Carlson-Stevermer, J. R. Zengel, C. M. Richards, P. Fozouni, J. Oki, L. Rodriguez, B. Joehnk, K. Walcott, K. Holden, A. Sil, J. E. Carette, N. J. Krogan, M. Ott, A. S. Puschnik, Genetic Screens Identify Host Factors for SARS-CoV-2 and Common Cold Coronaviruses. Cell. 184, 106–119.e14 (2021).

36. M. Varadi, S. Anyango, M. Deshpande, S. Nair, C. Natassia, G. Yordanova, D. Yuan, O. Stroe, G. Wood, A. Laydon, A. Žídek, T. Green, K. Tunyasuvunakool, S. Petersen, J. Jumper, E. Clancy, R. Green, A. Vora, M. Lutfi, M. Figurnov, A. Cowie, N. Hobbs, P. Kohli, G. Kleywegt, E. Birney, D. Hassabis, S. Velankar, AlphaFold Protein Structure Database: massively expanding the structural coverage of protein-sequence space with high-accuracy models. Nucleic Acids Res. 50, D439–D444 (2022).

37. J. Jumper, R. Evans, A. Pritzel, T. Green, M. Figurnov, O. Ronneberger, K. Tunyasuvunakool, R. Bates, A. Žídek, A. Potapenko, A. Bridgland, C. Meyer, S. A. A. Kohl, A. J. Ballard, A. Cowie, B. Romera-Paredes, S. Nikolov, R. Jain, J. Adler, T. Back, S. Petersen, D. Reiman, E. Clancy, M. Zielinski, M. Steinegger, M. Pacholska, T. Berghammer, S. Bodenstein, D. Silver, O. Vinyals, A. W. Senior, K. Kavukcuoglu, P. Kohli, D. Hassabis, Highly accurate protein structure prediction with AlphaFold. Nature. 596, 583–589 (2021).

38. P. Zhou, B. Li, Y. Yan, B. Jin, L. Wang, S. Y. Huang, Hierarchical Flexible Peptide Docking by Conformer Generation and Ensemble Docking of Peptides. J Chem Inf Model. 58, 1292–1302 (2018).

39. F. Chen, H. Liu, H. Sun, P. Pan, Y. Li, D. Li, T. Hou, Assessing the performance of the MM/PBSA and MM/GBSA methods. 6. Capability to predict protein-protein binding free energies and re-rank binding poses generated by protein-protein docking. Phys Chem Chem Phys. 18, 22129–39 (2016).

